# ‘*In plaque*-mass spectrometry imaging’ reveals a major metabolic shift towards odd-chain fatty acid lipids induced by host-virus interactions

**DOI:** 10.1101/317206

**Authors:** Guy Schleyer, Nir Shahaf, Carmit Ziv, Yonghui Dong, Roy A. Meoded, Eric J. N. Helfrich, Daniella Schatz, Shilo Rosenwasser, Ilana Rogachev, Asaph Aharoni, Jörn Piel, Assaf Vardi

## Abstract

Tapping into the metabolic cross-talk between a host and its virus can reveal unique strategies employed during infection. Viral infection is a dynamic process that generates an evolving metabolic landscape. Gaining a continuous view into the infection process is highly challenging and is limited by current metabolomics approaches, which typically measure the average of the entire population at various stages of infection. Here, we took a novel approach to study the metabolic basis of host-virus interactions between the bloom-forming alga *Emiliania huxleyi* and its specific virus. We combined a classical method in virology, plaque assay, with advanced mass spectrometry imaging (MSI), an approach we termed ‘*in plaque*-MSI’. Taking advantage of the spatial characteristics of the plaque, we mapped the metabolic landscape induced during infection in a high spatiotemporal resolution, unfolding the infection process in a continuous manner. Further unsupervised spatially-aware clustering, combined with known lipid biomarkers, revealed a systematic metabolic shift towards lipids containing the odd-chain fatty acid pentadecanoic acid (C15:0) induced during infection. Applying ‘*in plaque*-MSI’ might pave the way for the discovery of novel bioactive compounds that mediate the chemical arms race of host-virus interactions in diverse model systems.

## Introduction

Microbial interactions, such as those between a host and its pathogen, are shaped by a dynamic metabolic cross-talk. Gaining better understanding of this chemical communication has become an attractive approach to reveal basic metabolic principles that govern these interactions^1,2^. Metabolomics tools allow simultaneous identification of thousands of molecules in a given sample^3^, and have been used to monitor the metabolic fluctuations in diverse host-symbiont and host-pathogen model systems^4,5^.

Mapping the metabolic landscape of dynamic processes like viral infection requires high temporal resolution of the different stages of infection^5,6^. Most metabolic studies of viral infection are performed in bulk liquid cultures, measuring the average of the entire population at various time points during infection^7,8^. However, infection in liquid culture is not always a synchronous process, where all cells are infected at the same time. Asynchronous infection can lead to a mixed population of uninfected cells and infected cells at different stages of infection^9^. Consequently, bulk analysis might overlook rare subpopulations (e.g. resistant cells)^10^ and mask metabolic alterations that originate in these subpopulations, thus hindering our ability to track host defense mechanisms against infection^11^.

Rewiring of host metabolic pathways by the virus during infection generates multiple metabolic states which may affect the infection outcome (i.e. cell death or survival)^12^. Mapping these metabolic states into infection states might uncover unique strategies employed by the virus, which are essential for an optimal infection process^1^. Furthermore, occurrence of novel metabolic capabilities encoded by viruses (i.e. auxiliary metabolic genes), highly prevalence in the marine environment, provides a unique opportunity to uncover metabolic innovation during viral infection, shaped by host-virus co-evolution^13–15^. Recent advances in mass spectrometry imaging (MSI) technologies allow to localize specific metabolites of various dynamic processes and biotic interactions at the microscale level, and consequently to monitor metabolic changes in high spatiotemporal resolution^16,17^. MSI-based approaches were used to study seed germination^18^, metabolic footprints of host-associated bacteria^19,20^, embryonic development of zebrafish^21^, bacterial population dynamics^22–25^, to detect human pathogens^26^ and to visualize the spatial distribution of anti-viral drugs^27^.

An attractive host-virus model system is the cosmopolitan alga *Emiliania huxleyi* and its specific virus, *E. huxleyi* virus (EhV), which plays a key role in regulating the fate of carbon and sulfur cycles in contemporary oceans^28,29^. *E. huxleyi* forms massive annual blooms that cover vast oceanic areas and are terminated following infection by EhV^30,31^. EhV, a large dsDNA virus with a genome harboring 472 predicted protein coding sequences, encodes almost a full biosynthetic pathway for sphingolipids (SLs), a pathway never detected in any other known viral genome^32–34^. Virus-derived glycosphingolipids (vGSLs), products of this virus-encoded pathway, were found to be central components of the EhV membrane and to trigger host programmed cell death (PCD)^33–35^. vGSLs are produced exclusively during viral infection and are used as an effective metabolic biomarker to detect viral infection during *E. huxleyi* natural blooms in the ocean^33,36^. EhV infection causes a profound remodeling of *E. huxleyi*’s transcriptome, which leads to massive changes in the metabolome and lipidome of infected cells, for example changes in glycolysis, *de novo* fatty acid (FA) synthesis, downregulation of terpenoids pathways^35^ and induction of triacylglycerols (TAGs)^8,37,38^.

In this study, we sought to gain higher spatiotemporal resolution of metabolites that are associated with viral infection, and in addition, to discover new metabolites that are induced or reduced throughout this dynamic process. We combined a classical method in virology, plaque assay^39,40^, with advanced MSI technologies, a novel approach we termed ‘*in plaque*-MSI’. This approach increased the spatiotemporal resolution by which we could study the metabolic profile of viral infection, taking advantage of the characteristics of the plaque. MSI data were used to visualize the changes of known lipid biomarkers for viral infection across the plaque. In parallel, spatially-aware unsupervised clustering allowed a dedicated identification of lipids that were new to this host-virus model system as well as a potentially new class of lipids.

## Results

To gain high spatiotemporal resolution of the metabolic alterations during host-virus interactions, we mapped the metabolic profile of viral plaques (Figure 1). As opposed to infection in liquid medium, a plaque originates in a single infected cell^40^. The plaque expands via concentric rings, and each ring corresponds to the plaque circumference at a different time point, keeping a metabolic record of the infection^41,42^. Thus, a snapshot of a plaque unfolds the infection process in a continuous manner, from its initiation at the center to the uninfected area in its periphery. We used the virus-susceptible *E. huxleyi* CCMP2090 strain and the lytic virus EhV201 to perform plaque assays (Figure 1A). Plaques were visible to the naked eye usually two days post infection (dpi). Accurate identification of a plaque and its center prior to MSI was based on the algal host chlorophyll autofluorescence, as detected by epi-fluorescence microscopy^43^. The microscopic image served as a reference for the following MSI analysis. Two MSI techniques were used to analyze the plaques: matrix assisted laser desorption ionization-MS (MALDI, 60 μm resolution) and Flow-probe *in situ* micro-extraction^44^ (500 μm resolution) (Figure 1B). While MALDI-MS allowed identification of spatial patterns of metabolites at higher resolution, the Flow-probe-MS system had the advantage of faster sample analysis (~0.5 vs. ~10 hours per sample, respectively) and a matrix free preparation. The data collected by means of MSI was used to visualize the intensity of different metabolites along the plaque and to identify different spatial patterns (Figure 1C). The spatial dimension of the plaque revealed the potential link between changes of specific metabolites and infection dynamics.

**Figure 1:**
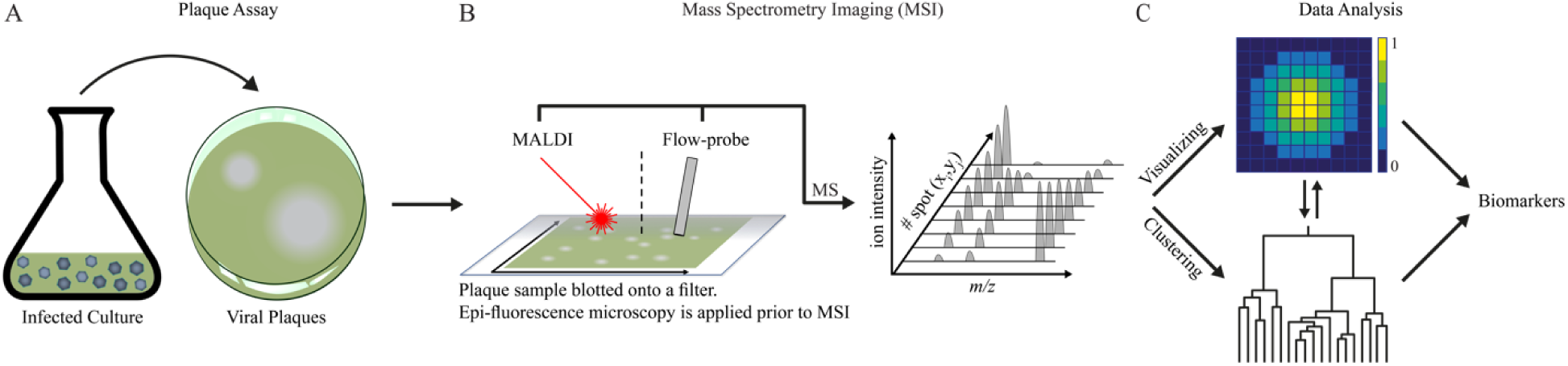
Overview of the workflow of ‘*in plaque*-MSI’ analysis. (A) Viral plaques were prepared by mixing infectious EhV virions with *E. huxleyi* host cells in agarose and pouring the mixture into a petri dish, which was then incubated until the formation of plaques. (B) A plaque sample was blotted onto a filter, which was imaged by epi-fluorescence microscopy prior to MSI in order to accurately identify the plaque morphology. Two MSI techniques were used to analyze plaque samples: MALDI-MS and Flow-probe-MS. (C) The collected spectra were used for visualization of specific metabolic markers for viral infection, as well as for spatially-aware unsupervised clustering, aiming at discovering novel metabolites associated with viral infection.

### Targeted MSI analysis of known lipid biomarkers across a viral plaque

We focused on known lipid biomarkers for viral infection, which were shown to be remodelled during infection in previous liquid chromatography-MS (LC-MS) based lipidomics experiments that were performed in liquid cultures^8,37,38^. These lipid biomarkers are usually classified based on their chemical category, such as: glycerolipids (GLs), glycerophospholipids (GPs) and sphingolipids (SLs). Since the viral genome also encodes the biosynthetic pathway for SLs^32,33^, the lipids can also be classified based on their biosynthetic origin (i.e. produced by alga- or virus-encoded enzymes). Additionally, these lipid biomarkers can be classified based on their relative abundance during infection (i.e. induction or reduction). Therefore, we grouped the lipids into three biological categories: virus-derived (encoded by the virus and induced during infection), host-induced and host-reduced (encoded by the host and induced or reduced during infection). vGSLs are the only virus-derived lipids known to date, with the majority of enzymes in their biosynthetic pathway encoded by the genome of the virus. vGSLs were found to be central components of the EhV membranes and to trigger host PCD during the lytic phase in a dose dependent manner^33–35^. Remodelled host lipids, on the other hand, include diverse classes: GLs such as BLL 36:6, TAG 48:1 and DGCC 40:7, and GPs such as PDPT 40:7 and PC 32:1, were all found to be induced during infection (i.e. ‘host-induced’), while others, such as BLL 38:6, were found to be reduced (i.e. ‘host-reduced’)^8,37,38^.

MS images of these lipid biomarkers across a plaque (Figure 2) allowed us to follow their induction and reduction patterns during infection in a continuous manner based on a snapshot view of the plaque. Despite the differences in resolution and sample preparation, both MALDI-MSI (Figure 2A) and Flow-probe-MSI (Figure 2B) presented comparable spatial patterns and highlighted the differences in the intensity of various lipids across the plaque. vGSL, for example, showed high intensity across the plaque area, while BLL 36:6 showed a different pattern, with lower intensity at the center of the plaque. Both lipids were detected only inside the plaque, whereas others, such as DGCC 40:7, PDPT 40:7 and TAG 48:1 appeared also outside the plaque area, albeit at lower intensity, indicating basal production in uninfected cells. Cross section profiles of each lipid (Figure 2A and B) revealed even finer differences, such as distance and degree of induction, between lipids with similar MS images. Induction of BLL 36:6, for example, was visible at a greater distance from the center than that of vGSL. Induction of DGCC 40:7 and PDPT 40:7 was visible at the same distance as BLL 36:6, however, their induction was more gradual than that of BLL 36:6 and vGSL. Interestingly, TAG 48:1 presented a unique ring-like pattern inside the plaque, not identified in other lipid species.

**Figure 2:**
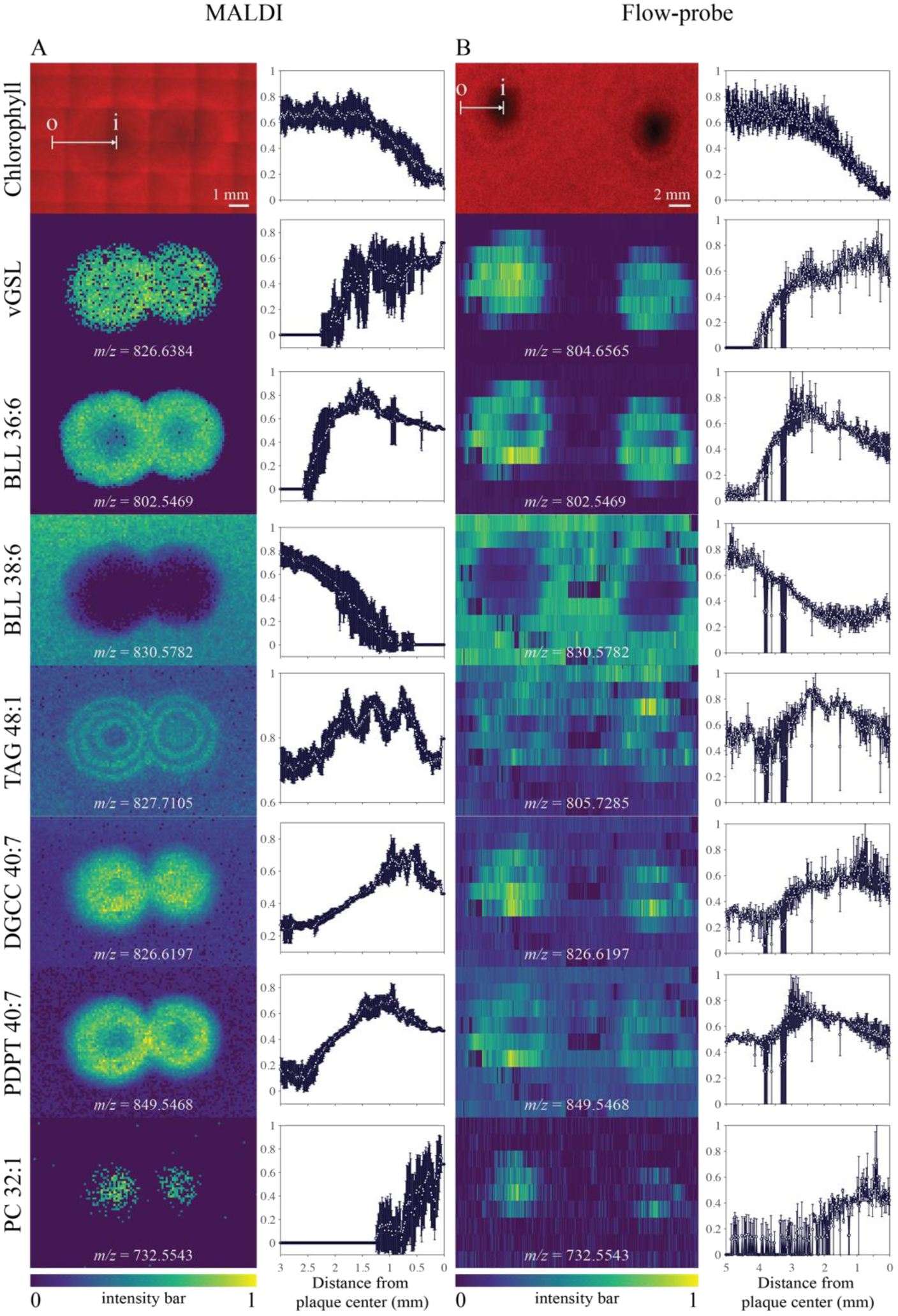
Targeted MSI analysis of known lipid biomarkers reveals the metabolic landscape produced during viral infection. Targeted analysis of two plaque samples at 5 dpi based on chlorophyll signal and seven lipid biomarkers for viral infection, using (A) MALDI-MS and (B) Flow-probe-MS. Chlorophyll autofluorescence (red pseudo-color) and MS imagesof specific lipids (false color representation of the detected ions) are presented. Cross section graphs show the relative intensity across the plaque, from its periphery to the center, as depicted by the arrows (‘O’, out; ‘I’, in). vGSL, viral glycosphyingolipid ([M+Na]^+^ = 826.6384 Da in MALDI-MS, [M+H]^+^ = 804.6565 Da in Flow-probe-MS); BLL 36:6, betaine-like lipid 36:6 ([M+H]^+^ = 802.5469 Da); BLL 38:6 ([M+H]^+^ = 830.5782 Da); TAG 48:1, triacylglycerol 48:1 ([M+Na]^+^ = 827.7105 Da in MALDI-MS, [M+H]^+^ = 805.7285 Da in Flow-probe-MS), DGCC 40:7, diacylglyceryl carboxyhydroxymethylcholine 40:7 ([M+H]^+^ = 826.6197 Da), PDPT 40:7, phosphatidyl-S,S-dimethylpropanethiol 40:7 ([M+H]^+^ = 849.5468 Da), PC 32:1, phosphatidylcholine 32:1 ([M+H]^+^ = 732.5543 Da)8,37,38. MS images were generated based on two samples (one for MALDI-MS and one for Flow-probe-MS). The analysis was performed on additional samples, which presented similar intensity patterns.

Taken together, the targeted analysis indicates the existence of chemotypic heterogeneity during infection, as visible from the diverse intensity patterns of specific lipids across the plaque. This continuous metabolic view of the infection process would not have been possible in bulk LC-MS-based lipidomics. The known lipid biomarkers and their respective intensity patterns further served as internal biomarkers for an untargeted analysis of the data.

### Spatially-aware clustering of Flow-probe-MS data

We applied an unsupervised spatially-aware clustering approach on Flow-probe-MS data generated from a plaque at 6 dpi in order to identify unknown metabolites that are altered during infection^45,46^. First, *m/z*-images were generated for each *m/z*-value recorded, depicting the intensity of the *m/z*-values in each of the pixels analyzed in the sample. The *m/z*-images were clustered based on correlations between their spatial intensity profiles (Figure 3A). A representative image was then generated for each cluster in the dendrogram, averaging the *m/z*-images of the cluster members. Based on these cluster-representative images, we selected clusters that presented distinct induction or reduction patterns across the plaque (Figure S1). Next, we matched the *m/z*-values in each cluster with the available annotations of lipid biomarkers (see Methods section for further details) and used them as markers for the cluster category (i.e. virus-derived, host-induced or host-reduced). For example, cluster two (CL2) included several vGSL species, and therefore was classified as a ‘virus-derived’ cluster, although it does not contain only virus-derived lipids. We then focused on four clusters which presented more pronounced changes across the plaque (Figure 3B). Three clusters (CL2, 4 and 5) were induced in the plaque and one (CL8) was reduced. In total, the four clusters contained 88 mass features (Table S1). Out of them, 62 mass features were putatively annotated as molecular ions, isotopes or fragments of lipids, corresponding with 36 feature groups, of which, 28 were induced and 8 reduced (Table S2). Fifteen feature groups were putatively annotated as lipids known from previous LC-MS-based studies (13 induced and 2 reduced)^8,37,38^. Sixteen were putatively annotated as lipids that are new to this host-virus model system (15 induced and 1 reduced, see Table 1 and Figure S2), one of which was putatively identified as vGSL-like t16:0/h22:0 (as previously suggested by long-chain base analysis^34^). Five were putatively identified as novel lipids (i.e. do not exist in our in-house repository or in public repositories) that share the same head group (all decreased, see Table 1 and Figures S2-S3), which we termed sulfonioglycerolipids.

**Figure 3:**
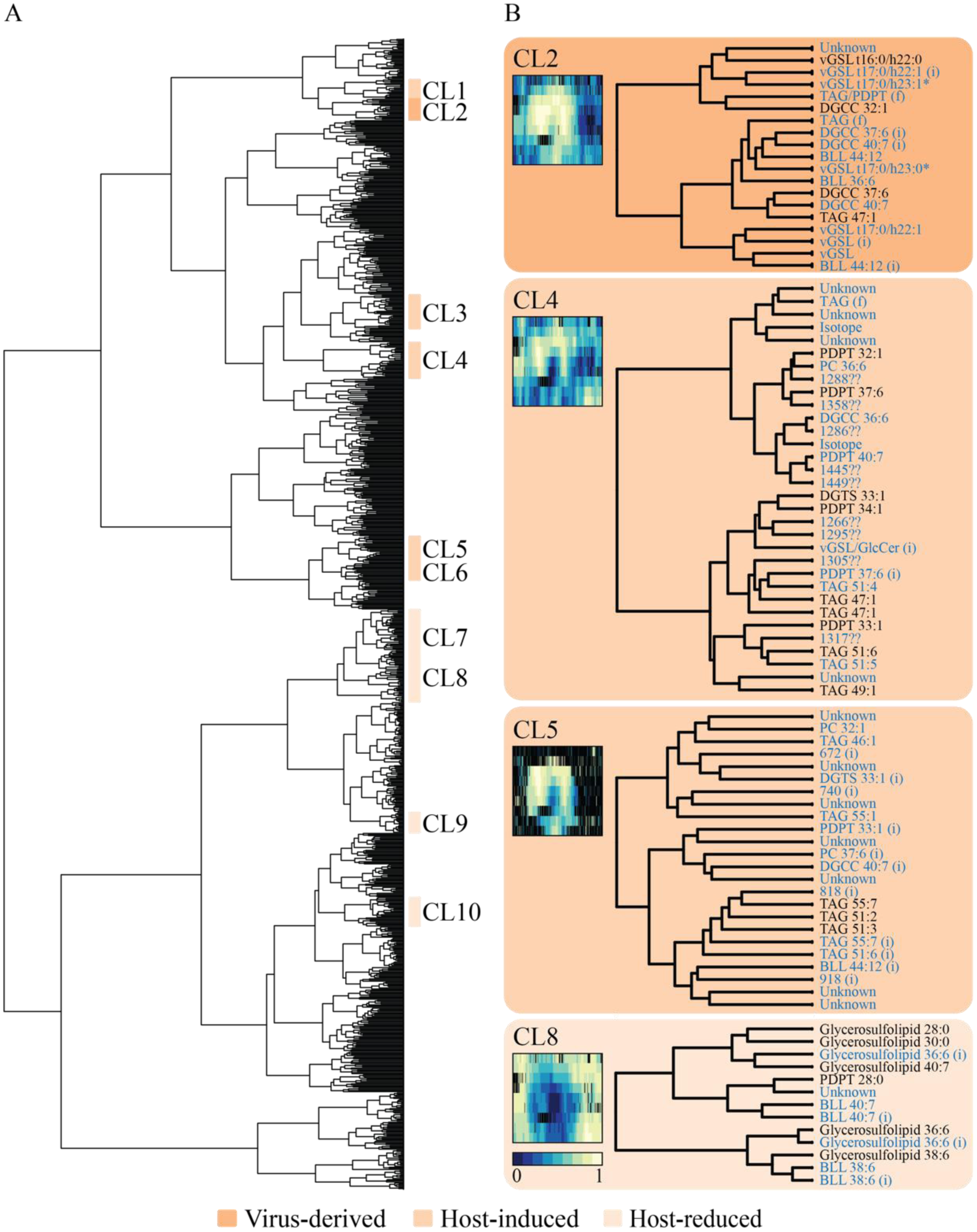
Unsupervised spatially-aware clustering of Flow-probe-MS data. (A) Dendrogram clustering of all *m/z*-values based on their spatial Flow-probe-MSI distribution along the plaque. Ten clusters that presented distinct induction or reduction patterns across the plaque (as was visible by the representative average images of the clusters, see Figure S1) were classified and labelled. The category of each cluster was determined by the inclusion of known lipid biomarkers for viral infection. (B) Four selected clusters are presented in detail, including their respective cluster-representative images and putatively annotated mass features. Mass features in black indicate lipids that are either new to *E. huxleyi*-EhV model system or belong to a potentially new class of lipids (Table 1). * mass features of vGSL t17:0/h23:1 and vGSL t17:0/h23:0 were also putatively annotated as GlcCer t18:0/h22:1 and GlcCer t18:0/h22:0, respectively. (i), isotope; (f), fragment.

**Table 1:** Newly identified lipids from four selected clusters and their putative identification.

| CL | Putative annotation<br>(lipid composition) | Theoretical<br>*m/z* | Chemical<br>formula | Change |
| --- | --- | --- | --- | --- |
| 2 | DGCC 32:1 (14:0/18:1) | 726.5884 | C<sub>42</sub>H<sub>77</sub>NO<sub>8</sub> | Induced |
| 2 | DGCC 37:6 (15:0/22:6) | 786.5884 | C<sub>47</sub>H<sub>79</sub>NO<sub>8</sub> | Induced |
| 2 | TAG 47:1 (14:0/15:0/18:1) | 791.7129 | C<sub>50</sub>H<sub>94</sub>O<sub>6</sub> | Induced |
| 2 | vGSL-like (t16:0/h22:0) | 790.6408 | C<sub>44</sub>H<sub>87</sub>NO<sub>10</sub> | Induced |
| 4 | DGTS 33:1 (15:0/18:1) | 724.609 | C<sub>43</sub>H<sub>81</sub>NO<sub>7</sub> | Induced |
| 4 | PDPT 32:1 (14:0/18:1) | 749.5155 | C<sub>40</sub>H<sub>77</sub>O<sub>8</sub>PS | Induced |
| 4 | PDPT 33:1 (15:0/18:1) | 763.5312 | C<sub>41</sub>H<sub>79</sub>O<sub>8</sub>PS | Induced |
| 4 | PDPT 34:1 (16:0/18:1) | 777.5468 | C<sub>42</sub>H<sub>81</sub>O<sub>8</sub>PS | Induced |
| 4 | PDPT 37:6 (15:0/22:6) | 809.5155 | C<sub>45</sub>H<sub>77</sub>O<sub>8</sub>PS | Induced |
| 4 | TAG 49:1 (15:0/16:0/18:1) | 819.7442 | C<sub>52</sub>H<sub>98</sub>O<sub>6</sub> | Induced |
| 4 | TAG 51:6 (18:0/18:1/18:5) | 837.6972 | C<sub>54</sub>H<sub>92</sub>O<sub>6</sub> | Induced |
| 5 | TAG 51:2 (15:0/18:1/18:1) | 845.7598 | C<sub>54</sub>H<sub>100</sub>O<sub>6</sub> | Induced |
| 5 | TAG 51:3 (15:0/18:1/18:2) | 843.7442 | C<sub>54</sub>H<sub>98</sub>O<sub>6</sub> | Induced |
| 5 | TAG 55:1 (15:0/18:1/22:0) | 903.8381 | C<sub>58</sub>H<sub>110</sub>O<sub>6</sub> | Induced |
| 5 | TAG 55:7 (15:0/18:1/22:6) | 891.7442 | C<sub>58</sub>H<sub>98</sub>O<sub>6</sub> | Induced |
| 8 | PDPT 28:0 (14:0/14:0) | 695.4686 | C<sub>36</sub>H<sub>71</sub>O<sub>8</sub>PS | Reduced |
| 8 | Sulfonioglycerolipid 28:0<br>(14:0/14:0)* | 689.5026 | C<sub>38</sub>H<sub>72</sub>O<sub>8</sub>S | Reduced |
| 8 | Sulfonioglycerolipid 30:0<br>(14:0/16:0)* | 717.5339 | C<sub>40</sub>H<sub>76</sub>O<sub>8</sub>S | Reduced |
| 8 | Sulfonioglycerolipid 36:6<br>(14:0/22:6)* | 789.5339 | C<sub>46</sub>H<sub>76</sub>O<sub>8</sub>S | Reduced |
| 8 | Sulfonioglycerolipid 38:6<br>(16:0/22:6)* | 817.5652 | C<sub>48</sub>H<sub>80</sub>O<sub>8</sub>S | Reduced |
| 8 | Sulfonioglycerolipid 40:7<br>(18:1/22:6)* | 843.5797 | C<sub>50</sub>H<sub>82</sub>O<sub>8</sub>S | Reduced |

Sixteen lipids are new to the E. huxleyi-EhV model system and five belong to a potentially new class of lipids (Sulfonioglycerolipids). The Lipids were putatively annotated based on MS/MS spectra (Metabolomics Standards Initiative level 2 annotation^47^), except those marked with an asterisk, which belong to the same putatively characterized compound class based on MS/MS spectra (level 3 annotation). CL, cluster (Figure 3). Change, as visible in the Flow-probe-MS images (Figure S2). [M+H]^+^ adduct is presented for all lipids. DGTS, Diacylglyceryl trimethylhomoserine.

Interestingly, 13 of the lipids that increased across the plaque were putatively identified as odd-chain fatty acid (OC-FA) lipids based on the Lipid Maps computationally-generated database of lipid classes and structure database (LMSD)^48^. Eleven were new to the *E. huxleyi*-EhV model system, while two were previously detected, however, their unique OC-FA characteristic was not recognized^38^. LC-MS-based analysis of these lipids during infection in liquid culture revealed profound induction as compared to control uninfected cells (Figure 4A and Table S3). These OC-FA lipids were also detected in LC-MS-based analysis of the plaques, as well as in purified virions and in extracellular vesicles that originated from infected cultures (Table S3).

**Figure 4:**
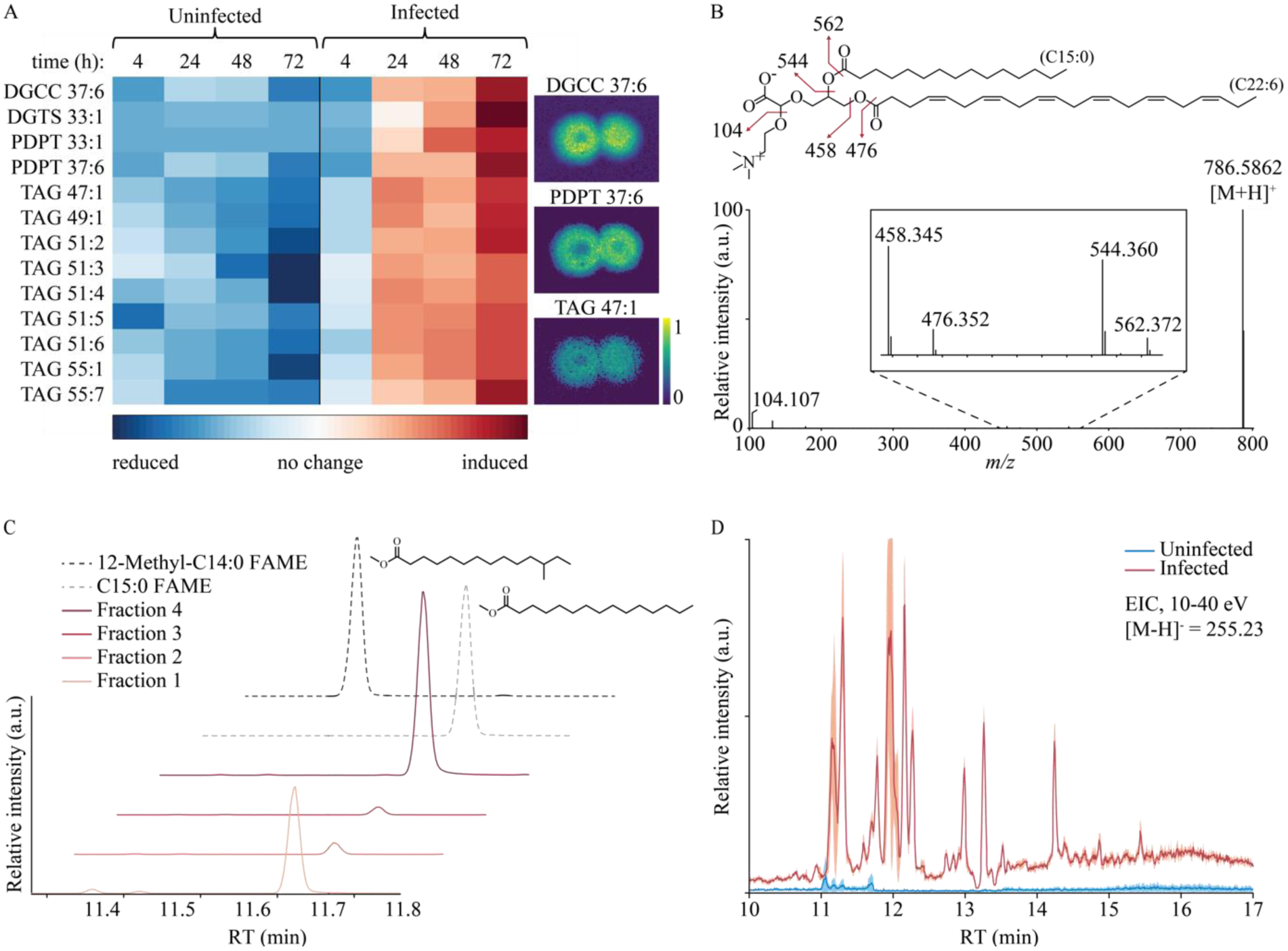
Induction of C15:0-based lipids during viral infection. (A) Heatmap representation of the alterations in C15:0-based lipids in uninfected and infected cells at different hours post infection (hpi) (left) and MALDI-MS images of three representative lipids (right). Differences between control and infected samples were significant for all lipid species (2-way ANOVA, fdr-corrected *P* < 0.02, *n* = 3). (B) LC-MS/MS putative identification of DGCC 37:6 with C15:0-FA (theoretical [M+H]^+^ = 786.5884 Da, observed = 786.5862 Da). Inset shows magnification of the mass range between 450-570 Da. Double bonds in the C22:6-FA were assigned based on the most common structure in the Lipid Maps structure database (LMSD). (C) GC-MS identification of C15:0 FAME in four different derivatized fractions collected from LC-MS runs of an infected culture at 48 hpi. The lipids collected in each fraction were derivatized by trans-esterification for FAME analysis. Fraction 1 – DGCC 37:6 and PDPT 37:6; Fraction 2 – PDPT 33:1; Fraction 3 – DGTS 33:1, Fraction 4 – TAGs. C15:0 FAME – Methyl pentadecanoate standard; 12-Methyl-C14:0 FAME – methyl 12-methylmyristate standard. (D) Systemic remodeling in C15:0-based lipids during viral infection presented by LC-MS extracted ion chromatogram (EIC) of C15:0 fragment ([M-H]^-^ = 255.23 Da, negative ion mode, energy ramp of 10-40 eV) in uninfected and infected cells after 48 h. Values are presented as mean ± s.d. (light red or blue color), *n* = 3.

MS/MS analyses of the OC-FA lipids using ‘*in plaque’*-Flow-probe revealed the existence of a C15:0-FA chain (pentadecanoic acid, Figure 4B and Table S2). The presence of C15:0-FA was further suggested by targeted LC-MS/MS analysis of infected cultures growing in liquid medium (Figures S4-S6 and Table S4). To validate the occurrence of C15:0-FA rather than a branched C14:0-FA (methyltetradecanoic acid), we performed hydrolysis and derivatization by trans-esterification on suspected lipids. The lipids were collected in four different fractions from an LC-MS run of an infected liquid culture, each fraction consisting at least one C15:0-based lipid (see Methods section for further details). The resulting fatty acid methyl ester (FAME) profiles were compared to analytical standards using gas chromatography-MS (GC-MS) analysis, which confirmed the existence of C15:0-FA chain (Figure 4C and Figure S7).

In order to explore the global distribution of C15:0-based lipids, we compared the abundance of C15:0-FA fragments in the lipidome of infected and uninfected cultures using LC-MS-based lipidomics of liquid cultures (Figure 4D). This comparison revealed massive induction of C15:0-based lipids during lytic infection in diverse lipid classes, suggesting a systemic-level metabolic switch towards a C15:0-based lipidome. Following a targeted search, we were able to putatively identify seven more lipids containing C15:0-FA (Table S5). One of these lipids was TAG 48:1, which was shown in previous works to be highly induced during infection^8^ (and across the plaque, where it presented a unique ring-like intensity pattern, see Figure 2), however, its FA composition was never explored. LC-MS/MS analyses indicated the existence of two structural isomers differing in their FA composition. While one isomer contained only even-chain FAs (C14:0, C16:0, C18:1) and was found both in uninfected in infected cells, the second isomer contained two OC-FAs (C15:0, C15:0, C18:1) and was found only in infected cells (Figure S8).

## Discussion

Tapping into the metabolic cross-talk between a host and its infecting virus can provide a new way for the identification of novel metabolic pathways that are essential for viral infection or host defence strategies. *E. huxleyi*-EhV is an important host-pathogen model system with great ecological significance in the marine environment^28,29,49,50^. Viral infection leads to substantial rewiring of the lipidome of infected cells, including a significant reduction in terpenoids^35^ and a profound induction in saturated TAGs^8^ and specific vGSLs^33,37,38^. The production of both TAGs and vGSLs is essential for the assembly of virions^8,34^. TAGs are also enriched in extracellular vesicles, which are produced in great numbers during infection and were shown to expedite infection across the population^51^. A major obstacle in studying host-virus interactions is to track this highly dynamic process and to map the metabolic landscape produced to a specific infection state. Gaining a continuous view into the infection process is limited by current metabolomics approaches, which typically measure the average of the entire population at various stages of infection. A viral plaque, on the other hand, captures the entire dynamics of the infection and resolves cells at different stages of infection in a spatiotemporal manner^41^.

By applying our MSI-based analyses to viral plaques, we were able to gain a high spatial resolution view of the metabolic continuum during infection. This ‘*in plaque*-MSI’ approach revealed massive induction of C15:0-based lipids during viral infection of *E. huxleyi* cells, as well as reduction in a potentially new class of lipids, termed sulfonioglycerolipids. This approach enabled us to harness the spatial structure of the plaque in order to detect differences in the induction and reduction patterns of known lipid biomarkers of viral infection, thus gaining high resolution of the dynamics of this process. These known lipids were shown to be remodelled during viral infection in previous LC-MS-based lipidomics experiments performed in liquid cultures and represented diverse lipid classes^8,37,38^. Rewiring of the host metabolome by the virus generates multiple metabolic states, as observed by host and viral gene expression^12^. In the future, the spatial structure of the plaque might allow us to distinguish between the different metabolic states along the plaque and use them to predict the infection outcome.

We could identify differences between lipids that previously presented similar trends during infection. vGSL and BLL 36:6, which are used as lipid biomarkers for *E. huxleyi* infection in the natural environment^33,36,52^, were both induced in the plaque. However, BLL 36:6 was reduced at the center of the plaque and was induced at a greater distance from the center than vGSL (Figure 2). Lower intensity of BLL 36:6 at the center might suggest modification of this lipid or consumption and recycling to other metabolites at later stages of infection. The different distance of induction might be explained by production of BLL 36:6 in cells at earlier stages of infection than vGSL, or even in uninfected cells due to a chemical signal diffusing from the infected cells. It is also possible that BLL diffuses at a faster rate than vGSL in the plaque. Our findings suggest that BLL 36:6 may be used as a biomarker for earlier stages of infection, which are otherwise not detected when using vGSL alone. Presence of both vGSL and BLL 36:6, on the other hand, might be indicative of a population at later stages of infection, prior to cell lysis.

The structure of the plaque and the identification of distinct induction and reduction patterns across the plaque allowed us to apply unsupervised spatially-aware clustering on Flow-probe-MS data in order to identify novel metabolites that are induced or reduced during infection. The representative average image of each cluster was used to reduce the complexity of the untargeted data, allowing us to select clusters with biological relevance to the infection dynamics. Identification of known lipid biomarkers for infection in each of these clusters (e.g. vGSL in cluster 2) was used to define their category (virus-derived, host-inducted and host-reduced). A selected number of clusters, all of them with distinct patterns, enabled us to concentrate our identification efforts on a smaller number of mass features, further reducing the complexity of the analysis (Figure 3). Thus, we were able to detect a group of functionally relevant metabolites from a plethora of unknown mass features based on their spatial intensity patterns. Such dedicated identification might help to uncover metabolites that are masked in untargeted LC-MS-based metabolomics analyses of liquid cultures. Applying *‘in plaque*-MSI’ to other host-virus interactions might allow the discovery of novel bioactive molecules and potential anti-viral compounds.

We putatively annotated 16 lipid species that were new to this host-virus model system and five of a potentially new class of lipids. More than half of the lipids were putatively identified as C15:0-based lipids and were highly increased across the plaque and in infected liquid cultures (Figure 4). Interestingly, C15:0-FAs were incorporated in lipids usually found in the cytoplasmic membrane (e.g. SL, DGCC, DGTS and PDPT) or in lipid droplets (e.g. TAG), but not in lipids found in the chloroplast (e.g. Glycosyldiradylglycerols)^53^. While the biological function of OC-FA lipids is still unknown, C15:0-FA has been previously shown to be less preferred for β-oxidation than even-numbered FAs^54,55^. C15:0-FA is also involved in the biosynthesis of vGSL. Viral infection causes a shift in the substrate specificity of the enzyme pyridoxal 5′-phosphate (PLP)-dependent serine palmitoyl transferase (SPT; EC 2.3.1.50), the rate limiting enzyme of the entire SL biosynthetic pathway. In infected cells, the SPT enzyme was shown to use C15:0-FA-CoA, while it typically utilizes a C16:0-FA-CoA in uninfected cells^34^. The C15:0-FA-CoA used by viral SPT serves as a substrate for the unusual hydroxylated C17 long-chain base found in the vGSLs^34^. We further observed a profound systematic induction in C15:0-based lipids, as detected by LC-MS analysis of liquid cultures (Figure 4). Selective incorporation of OC-FA lipids that are less prone to β-oxidation^54,55^ in the membrane might increase the durability of the viral particles and vesicles, and consequently attenuate their decay rate in the marine environment. Induction of C15:0-based lipids during viral infection is also consistent with a previous report that indicated an increase in the levels of C15:0-FA during viral infection, although the specific lipids were not investigated^56^.

Recent years have witnessed a growing interest in lipids containing OC-FA and their role as markers for metabolic disorders^57–60^. OC-FA lipids, including C15:0-based lipids, have been reported in humans, animals, microorganisms and plants^61–66^. Nevertheless, understanding of their biological function is still elusive. Several biosynthetic pathways have been proposed for OC-FAs, either *de novo* or by chain-shortening of longer FA chains. *De novo* synthesis involves the conversion of propionic acid to propionyl-CoA, which, in turn, can replace acetyl-CoA in the initial step of fatty acid synthesis^67,68^. Excess of propionyl-CoA was reported to increase the levels of C15- and C17-FAs in humans^57,69^. In addition, phytosphingosine, a sphingoid base of GSLs, was recently reported as a source of C15:0-FAs in yeast. The odd FAs are later incorporated into GPs^70^. Future studies will enable elucidation of the biochemical origin and function of C15:0-FA lipids during viral infection in the ocean.

Taken together, our work revealed a systematic metabolic shift towards C15:0-based lipids as a result of viral infection. This shift might be part of the viral strategy to hijack host metabolism during infection, by inducing an orthogonal biosynthetic pathway for the synthesis of these lipids.

Given their high and specific abundance during infection, C15:0-based lipids might serve as novel biomarkers for *E. huxleyi*-EhV interactions in the ocean.

## Methods

### Culture growth and viral infection dynamics

*E. huxleyi* strain CCMP2090 was used for this study. Cells were cultured in K/2 medium^71^ in artificial seawater (ASW)^72^ supplemented with ampicillin (100 μg mL^-1^) and kanamycin (50 μg mL^-1^), and incubated at 18 °C with a 16:8 h light:dark illumination cycle. A light intensity of 100 μmol photons m^-2^ s^-1^ was provided by cool white light-emitting diode lights. The virus used for this study is the lytic *E. huxleyi* virus EhV201^73^.

### Enumeration of cells and viruses

Cells were quantified using a Multisizer 4 Coulter Counter (Beckman Coulter, version 4.01) and an Eclipse (iCyt) flow cytometer (Sony Biotechnology, Champaign, IL, USA, using ec800 version 1.3.7 software), equipped with 405 and 488 nm solid state-air cooled lasers (both 25 mW on the flowCell) and standard filter setup. Algae were identified by plotting chlorophyll fluorescence in the red channel (737 to 663 nm) versus green fluorescence (500 to 550 nm) or side scatter. Extracellular viral particles were quantified as described previously^34^.

### Plaque assay

150 mL of cells at 2-3×106 cells mL^-1^ were concentrated (2000 ×g, 5 min, 18 °C) to 2.7 mL. 300 μL of virions at a concentration of 10^4^ virions mL^-1^ were added to the cells. After 2 h of incubation under normal growth conditions, the host-virus mixture was mixed with 9 mL of K/2 medium in ASW that contained 0.2% agarose (SeaKem LE agarose, Rockland, ME, USA) and then poured onto an agarose plate (12×12 cm) containing K/2 medium in ASW solidified by a 1.5%, supplemented with ampicillin (100 μg mL^-1^) and kanamycin (50 μg mL^-1^). Plaques were visible to the naked eye usually at 2 dpi. Plaque samples were subjected to MSI analysis at 5-6 dpi, when they were at a size large enough for spatial analysis (diameter of 4-8 mm) yet were still active and growing. Blank samples were prepared following the same procedure, using K/2 medium only (i.e. without the host-virus mixture).

### Sample preparation for epi-fluorescence microscopy and MSI

Plaque samples were cut using a metal square punch and transferred to a glass slide. The upper layer of 0.2% agarose was blotted onto a 0.22 μm PVDF filter (Merck Millipore, Cork, Ireland). The filter containing the plaque was then dried for ~10 min at 30 °C. The filter was immobilized on a microscope slide with double-sided adhesive tape for further analyses using epi-fluorescence microscopy and MSI.

### Epi-fluorescence microscopy

Microscopy images used for both Flow-probe-MS and MALDI-MS samples where obtained using Olympus IX81 motorized epi-fluorescence inverted optical microscope (Olympus, Tokyo, Japan) equipped with ×4 (numerical aperture (NA) 0.13, for MALDI-MS samples) and ×10 (NA 0.3, for Flow-probe-MS samples) objectives and a filter system for chlorophyll autofluorescence (Flow-probe-MS samples: ex: 470/40 nm, em: 632/60 nm, MALDI-MS samples: ex: 430/24 nm, em: 590 nm LP). Images were captured using ORCA-Flash 4.0 V2 (Hamamatsu Photonics KK, Hamamatsu City, Japan, for Flow-probe-MS samples) or Coolsnap HQ2 CCD (Photometrics, Tuscon, AZ, USA, for MALDI-MS samples) cameras and processed using MetaMorph (version 7.8.7, Molecular Devices) and CellSens Dimension (version 1.11, Olympus) software packages.

Image analysis and processing was done using Fiji (Fiji Is Just ImageJ, version 1.51w). To correct for uneven illumination and remove background, a background image was generated by projecting the median of all frames and performing Gaussian blur (30 pixels diameter). Then, all frames were divided by the background image. Frames were then stitched with 10% tile overlap. The microscopy images allowed accurate identification of a plaque and its center prior to MSI. The center of the plaque can be detected by a reduced chlorophyll signal and is consisting of mainly lysed cells (Figure 2)^43^. The infected area surrounding the center, on the other hand, is populated by living cells, as indicated by the chlorophyll signal.

### Flow-probe-MS imaging

The analysis was performed at ambient conditions using a ‘Flow-probe’ *in situ* micro-extraction system^44^ (Prosolia, Indianapolis, IN, USA), connected to a Thermo Scientific ‘Qexactive’ mass spectrometer. The ‘Flow-probe’ was operated using Flowprobe nMotion software (version 1.0.0.58, Prosolia). The instrument was operated in positive ion mode and the spray voltage set to 3 kV with a capillary temperature of 150 °C. A mixture of methanol:acetonitrile:toluene 50:35:15 + 0.1% TFA^74^ with a flow-rate of 10 μL min^-1^ was used. Samples were analyzed by measuring several parallel lanes (one mm wide, spacing one mm) using a probe-speed of 100 μm sec^-1^. Chemical entities were monitored in a mass range between 125 and 1100 *m/z* at a resolution of 70,000 *m/z* for MS1 scans. Data-dependent fragmentation was performed on ten precursor ions per MS1 scan at a resolution of 17,500 *m/*z. Simultaneous Flow-probe imaging and data-dependent acquisition were performed either using an inclusion list of known lipids, while allowing ions not included in the inclusion list to be fragmented in the absence of said ions or without an inclusion list and a dynamic inclusion window of three seconds. Full list of the lipids annotated using Flow-probe-MS/MS and their fragments can be found in Table S2.

### Processing of Flow-probe-MSI data

Raw Flow-probe-MSI data was converted from the vendor’s format to the open format mzML using ‘msconvert’ software (part of ProteoWizard version 3.0)^75^ and from the mzML format to the MSI compatible data format imzML^76^ using ‘imzMLConverter’ software (version 1.3)^77^. Next, imzML files corresponding with each experimental run were imported into the R open source programming environment (www.r-project.org) using the R package ‘MALDIquant’^78^. Initially, 75 heading and trailing scans were removed from each horizontal acquisition line due to high levels of noise and measurement calibration issues, and one horizontal acquisition line (out of ten) was completely removed from analysis due to low observed performance of the instrument. Low intensity scans, with a total ion current (TIC) lower than the 7.5% quantile of the overall scan TIC statistics (203 scans in total) were like-wise removed immediately after raw data import and before subsequent peak detection steps, to reduce biases or shifts in calculation of mass measurement values. Next, software parameters for mass peak detection, smoothing, alignment, binning and filtering were determined using manual inspection of mass peaks in the raw data and according to the instrument’s mass measurement specifications, as suggested in the ‘MALDIquant’ software guidelines (see Table S6). The initial preprocessing step after quality control scan filtering procedure resulted in a scan to mass feature intensity matrix of 7035 mass features measured across an average of 428.5 scans per flow-probe horizontal acquisition line (corresponding to a matrix dimensions of 3856 scans by 7035 mass features).

Next, matrix intensity values were log transformed and normalized using a series of 25 masses detected in separate injections of a ‘blank’ material (i.e. Flow-probe-MS analysis of a culture-free sample derived from K/2 medium in ASW containing 0.2% agarose) which were used as baseline reference values for the analytical measurement variance, independent of biological factors. Thus, the median of measured intensity values corresponding with the detected ‘blank’ masses was subtracted from the measured intensity values of all mass features across all scans, resulting in a feature matrix of log transformed and analytically-normalized arbitrary intensity values in the range of -6.46 to 4.91. The feature matrix was finally reduced column-wise by removal of sparse mass features which contained more than 99% missing values (5295 features, 75.26% of all mass features). The remaining 1740 mass features were then used to compute an intensity profile correlation matrix, using Spearman’s rho measure of association between all complete (non-NA) pairs of intensity profiles, resulting in a 1740 by 1740 correlation matrix. Agglomerative hierarchical clustering was then performed by transforming the correlation matrix into a distance matrix (with values in the range zero to one) using the ‘agnes’ function in the R package ‘cluster’ [“*Maechler, M., Rousseeuw, P., Struyf, A., Hubert, M., Hornik, K.(2016). cluster: Cluster Analysis Basics and Extensions. R package version 2.0.5.*”]. Further cluster analysis was performed by converting the cluster to a dendrogram object and focusing on clusters of mass profiles related to annotated masses of known (i.e. previously identified) biologically meaningful lipids or by visual inspection of cluster images containing spatial morphologies. Annotation of mass features to masses of known lipids was done using an in-house R script which performed mass-to-mass comparisons of observed mass features with the theoretical accurate mass of the protonated ([M+H]^+^), sodium ([M+Na]^+^) and ammonium ([M+NH_4_]^+^) ions of each known lipid, using a mass error tolerance of 6 parts per million (ppm). The annotation script further extracted the natural heavy isotope pattern of each putatively annotated parent ion and performed chemical formula decomposition using the ‘decomposeMass’ in the R package ‘Rdisop’79. Putative annotations of mass features to known lipids were then assigned using the combined criteria of: mass-to-mass, chemical formula decomposition and manual inspection of the corresponding mass peak in the raw data. Automatic generation of cluster spatial morphology images to aid manual cluster inspection was performed using an in-house R script which transformed the intensity profiles of each sub-cluster to an image matrix of 9 by 450 pixels and which plotted the mean sub-cluster intensity as an image. A minimum threshold of five mass features per sub-cluster was used for recursive bottom-up traversing of the cluster dendrogram and plotting was performed using the ‘YlGnBu’ in the R package ‘RColorBrewer’ [“*Erich Neuwirth (2014). RColorBrewer: ColorBrewer Palettes. R package version 1.1-2*”].

MS-images of specific *m/z* values were generated using the open-source software package MSiReader v1.00^80,81^, built on the platform of Matlab (Mathworks, Natick, MA, USA) using ‘Viridis’ colormap.

### Matrix sublimation for MALDI-MSI

2,5-Dihydroxybenzoic acid (DHB) matrix was deposited using a sublimation apparatus (height × inner diameter, 250 mm × 152 mm, Sigma-Aldrich, St Louis, MO, USA). The sublimator was coupled to a rough pump and a digital thermocouple vacuum gauge controller and was placed on a sand bath heated by a hot plate. The temperature was monitored by a digital thermometer. 200 mg of DHB matrix were sublimated under a fixed pressure of 8×10^-2^ Torr at 140 °C for 4 min.

### MALDI-MS imaging and data processing

MALDI imaging experiments were performed using a 7T Solarix FT-ICR (Fourier transform ion cyclotron resonance) mass spectrometer (Bruker Daltonics, Bremen, Germany). Datasets were collected in positive mode using lock mass calibration (DHB matrix peak: [3DHB+H-3H2O]^+^, *m/z* 409.055408) in the range of 150-3000 *m/z,* with a spatial resolution of 60-100 μm. Each mass spectrum was obtained from a single scan of 100 laser shots at a frequency of 1 kHz and a laser power of 18%. The acquired spectra were processed using FlexImaging software (version 4.0, Bruker Daltonics, Bremen, Germany) and SCiLS Lab 2015b (SCiLS GmbH, Bremen, Germany). Data sets were normalized to RMS intensity and MALDI images were plotted at the theoretical *m/z* ± 0.001% (FlexImaging) and *m/z ±* 5 ppm (SCiLS Lab), with pixel interpolation on. MS-images were produced using SCiLS Lab 2015b using ‘Viridis’ colormap.

### Cross section graphs

Cross section graphs were plotted by averaging the intensity of each lipid or chlorophyll signal across several lines from the outer part of the plaque to its center. The center was selected to be the minimum of chlorophyll signal. Ten lines were used in fluorescent microscopy and MALDI-MS and two in Flow-Probe-MS images. Intensity values for each line were extracted from the images using Fiji. Graphs were plotted using Matlab R2016b.

### Chemicals and internal standards for LC-MS analyses

Liquid chromatography-grade solvents were purchased from Merck (Darmstadt, Germany) and Bio-Lab (Jerusalem, Israel). Ammonium acetate was purchased from Sigma-Aldrich (St Louis, MO, USA). Internal standards for lipidomics analysis were purchased from Avanti Polar Lipids (Alabaster, AL, USA). The lipid internal standards were added to the extraction solution employed in the initial extraction step.

### Lipidomics Analyses

Lipids were extracted from liquid cultures of *E. huxleyi* cells infected with EhV201 and from non-infected cells harvested at 4, 24, 48 and 72 hpi in three biological replicates. Infection was performed on exponential phase cultures (5×10^5^ to 10^6^ cells mL^-1^), which were infected 2-3 h after the onset of the light period, with a 1:100 volumetric ratio of viral lysate to culture (multiplicity of infection of 1:1 viral particles per cell). The samples (30-150 mL of each culture, equivalent to ~5×10^7^ cells per sample) were collected on GF/C filters (pre-combusted at 400 °C for 8 h), plunged into liquid nitrogen, and stored at -80 °C until analysis^33^.

Lipid extraction and analysis were performed as previously described^82^ with some modifications: filters containing algae were placed in 15 mL glass tube and extracted with 3 mL of a pre-cooled (-20 °C) homogenous methanol: methyl-tert-butyl-ether (MTBE) 1:3 (v/v) mixture containing 0.1 μg mL^-1^ of glucosylceramide (d18:1/12:0) which was used as an internal standard. The tubes were shaken for 30 min at 4 °C and then sonicated for 30 min. Ultra-performance liquid chromatography (UPLC) grade water:methanol (3:1, v/v) solution (1.5 mL) was added to the tubes followed by centrifugation. The upper organic phase (1.2 mL) was transferred into 2 mL Eppendorf tube. The polar phase containing algae debris and filter pieces was re-extracted with 0.5 mL of MTBE. Organic phases were combined and dried under N_2_ stream and then stored at -80 °C until analysis. Extraction of lipids from plaque samples was performed by suspending the upper soft layer containing the plaque in the lipid extraction solution, following the same procedure described above (using third of the mentioned volumes). As a blank, soft layer without *E. huxleyi* cells (i.e. ASW + K/2 containing 0.2% agarose only) was extracted as well. Samples were extracted in three replicates. Isolation, concentration and lipid extraction of vesicles and virions was performed as previously described^51^. All dried lipid extracts were re-suspended in 300 μL buffer B (see below) and centrifuged again at 18,000 ×g and 4 °C for 5 min.

### UPLC-q-TOF MS for lipidomics analysis

Post extraction, the supernatant was transferred to an autosampler vial and an aliquot of 1 μL was subjected to UPLC-MS analysis. Lipid extracts were analyzed using a Waters ACQUITY UPLC system coupled to a SYNAPT G2 HDMS mass spectrometer (Waters Corp., Milford, MA, USA). Chromatographic conditions were as described previously^82^. Briefly, the chromatographic separation was performed on an ACQUITY UPLC BEH C8 column (2.1×100 mm, i.d., 1.7 μm) (Waters Corp., Milford, MA, USA). The mobile phase consisted of water (UPLC grade) with 1% 1 M NH_4_Ac, 0.1% acetic acid (mobile phase A), and acetonitrile:isopropanol (7:3) with 1% 1 M NH4Ac, 0.1% acetic acid (mobile phase B). The column was maintained at 40 °C and the flow rate of the mobile phase was 0.4 mL min^-1^. MS parameters were as follows: the source and de-solvation temperatures were maintained at 120 °C and 450 °C, respectively. The capillary voltage and cone voltage were set to 1.0 kV and 40 V, respectively. Nitrogen was used as de-solvation gas and cone gas at the flow rate of 800 L h^-1^ and 20 L h^-1^, respectively. The mass spectrometer was operated in full scan MS^E^ positive resolution mode over a mass range of 50-1500 Da. For the high energy scan function, a collision energy ramp of 15-35 eV was applied, for the low energy scan function -4 eV was applied. Leucine-enkephalin was used as lock-mass reference standard. The major ions and specific fragment ions of the lipids were analyzed in positive ionization mode. For detection of C15:0-FA fragment ions of the lipids, samples were injected also in negative ionization mode. The MS^E^ energies applied for the negative ionization were 15-40 eV. For MS/MS (in positive ionization mode), collision energy ramp of 10-45 eV was applied.

### LC-MS and LC-MS/MS data analysis, lipid identification and quantification

LC-MS data were analyzed and processed with MassLynx and QuanLynx (version 4.1, Waters Corp., Milford, MA, USA). The putative identification of the various lipid species was performed according to the lipid accurate mass and LC-MS/MS fragmentation pattern, as described previously^34,37,38^. The putative identification of lipid species that were new to this host-virus model system (listed in Table 1) was based on the Lipid Maps computationally-generated database of lipid classes and structure database (LMSD)^48^ (Metabolomics Standards Initiative level 2 annotation47). See Figures S4-S6 for MS/MS spectra and list of fragments of representative species of each class. Five unidentified mass features, later termed sulfonioglycerolipids (listed in Table 1), did not exist in our in-house repository or in public repositories such as Lipid Maps^48^ and METLIN^83^. Putative structures of these sulfonioglycerolipids, which belong to the same putatively characterized compound class (level 3 annotation), was based on accurate mass and LC-MS/MS fragmentation pattern, as elaborated in Figure S3. Full list of the lipids annotated using LC-MS/MS and their fragments can be found in Table S4 and Table S5. Relative levels of lipids extracted from liquid cultures were normalized to the internal standard and the number of algal cells used for analysis. Heatmap was generated using Matlab R2016b. Mean intensity values (normalized peak intensity per cell, *n* = 3) were log_10_ transformed and standardized per lipid (row). Zero values were replaced with 0.5 of the minimal value prior to log_10_ transformation.

### Fatty acid composition

Four fractions enriched with potential C15:0-based lipids were collected manually from LC-MS runs of lipidomics extracts from infected cells at 48 hpi. Fractions were collected at the following RTs: fraction 1, 4.3-5.3 (containing DGCC 37:6 and PDPT 37:6); fraction 2, 6.6-7.4 (containing PDPT 33:1); fraction 3, 7.5-8.4 (containing DGTS 33:1); fraction 4, 16.1-18.3 (containing TAGs). Presence of the above-mentioned lipids in the collected fractions was verified by re-injecting the collected fractions to the LC-MS, using the same conditions for lipidomics analysis. The collected samples were dried under N_2_ flow. As a negative control, 4 mL of mobile phase A and 4 mL of mobile phase B were dried as well. As a positive control, 6 mL of the extraction solution for lipids containing isotopically labelled C15:0-FA containing TAG 48:1 (C15:0/C18:1/C15:0) standard (Avanti Polar Lipids, Alabaster, AL, USA) were dried as well.

The FA composition of lipids was analyzed using derivatized fatty acid methyl esters (FAMEs). The samples were dissolved in 500 μL 1.25 M methanolic hydrochloride (MeOH/HCl, Sigma-Aldrich, St. Louis, MO, USA), incubated for 1 h at 80 °C while shaking, and then dried under N_2_ flow. The resulting FAMEs were extracted twice with 250 μL hexane:chloroform 1:1 and once with 250 μL chloroform, each extraction for 30 min at room temperature while shaking. The extracts were dried under N_2_ flow. The dried FAME extracts were re-suspended in 100 μL hexane and centrifuged at 18,000 ×g and 4 °C for 5 min.

### GC-MS for FAME analysis

Post extraction, the supernatant was transferred to an autosampler vial and an aliquot of 1 μL was subjected to GC-MS. The GC-MS system comprised an Agilent 7890A gas chromatograph equipped with split/splitless injector, and LECO Pegasus HT Time-of-Flight Mass Spectrometer (TOF-MS, LECO Corp., St Joseph, MI, USA). GC was performed on a 30 m × 0.25 mm × 0.25 μm Rxi-5Sil MS column (Restek). Samples were analyzed in the splitless mode; injector temperature was set at 280 °C. Analytes by 1 μL injected were separated using the following chromatographic conditions: helium was used as carrier gas at a flow rate of 1.0 mL min^-1^. The thermal gradient started at 80 °C and was held at this temperature for 2 min, ramped to 330 °C at 15 °C min^-1^ and then held at 330 °C for 6.0 min. Eluents were fragmented in the electron impact mode with an ionization voltage of 70 eV. The MS mass range was 45-800 *m/z* with an acquisition rate of 20 spectra per second. The ion source chamber was set to 250 °C and the transfer line to 250 °C. LECO ChromaTOF software (version 4.50.8.0, LECO Corp.) was used for acquisition control and data processing. FAMEs were identified by comparison of their mass spectra and retention times to the corresponding standards, injected in the same GC-MS conditions. Two standards were used: methyl pentadecanoate (C15:0 FAME, Sigma, St. Louis, MO, USA) and methyl 12-methylmyristate (Methyl 12-C14:0 FAME, Santa Cruz Biotechnology, CA, USA), at a concentration of 10 μg mL^-1^ in hexane.

## Data availability

Data supporting the findings of this study are available within the paper (and its supplementary information files). Raw data generated or analyzed during the current study will be available in the future.

## Acknowledgements

We thank Constanze Kuhlisch from the Vardi lab for her assistance with GC-MS analyses and fruitful discussions, Shiri Graff van Creveld from the Vardi lab for her assistance in designing the figures for this manuscript, and Avia Mizrachi from the Vardi lab for her assistance with image analysis and processing. We also thank Alexander Brandis from the Targeted Metabolomics Unit at the Life Sciences Core Facilities, Weizmann Institute of Science, for his assistance in FAME derivatization, Ron Rotkopf from the Bioinformatics Unit, Department of Biological Services, Weizmann Institute of Science, for his assistance with the statistical analysis, Sung Sik Lee from the scientific Center for Optical and Electron Microscopy (ScopeM), ETH Zürich for his assistance with epi-fluorescence microscopy, and Tal Luzzatto-Knaan from the Department of Marine Biology, University of Haifa, for her useful comments on the manuscript. This research was supported by the European Research Council CoG (VIROCELLSPHERE grant no. 681715) awarded to A.V and by EMBO Short Term Fellowship (ASTF 601 - 2015) awarded to G.S.

## Author contributions

G.S. and A.V. conceptualized the project and conceived and designed the experiments, G.S. and A.V. wrote the manuscript; G.S. performed all experiments; N.S. developed computational analysis of MS-data.; C.Z. conducted lipid extractions and LC-MS experiments; R.A.M and E.J.H.E conducted Flow-probe-MS experiments; Y.D. conducted MALDI-MS experiments; I.R. conducted GC-MS experiments; D.S. isolated vesicles and virions for lipidomics analysis; all authors provided useful feedback on the experimental design and comments on the manuscript.

